# The pH gradient contributes to persistence in *Mycobacterium tuberculosis*

**DOI:** 10.1101/2025.09.12.675885

**Authors:** Hassan E. Eldesouky, Kristin N. Adams, Justin K. Brache, Laarni Kendra T. Aguila, Mariana Garcia, Enming Xing, Pui-Kai Li, David R. Sherman

## Abstract

Tuberculosis (TB) remains difficult to cure due in part to poorly defined drug-tolerant persister cells formed by *Mycobacterium tuberculosis* (Mtb), which survive antibiotic treatment without evidence of genetic resistance. To better define this phenotype, we screened 2,336 FDA-approved drugs for compounds that target persistence. Unexpectedly, we identified a strong inducer of drug tolerance -- the antiparasitic niclosamide (NCA), which is known to disrupt proton motive force. In contrast to earlier reports that it harbors promising anti-TB activity, we found that NCA protected Mtb from bactericidal doses of isoniazid, rifampicin, and other standard TB drugs. Investigating further, we showed that disruption of the pH gradient and consequent intracellular acidification is needed to induce tolerance, while disruption of membrane potential is not, and also that protection is tunable by external pH. Transcriptomic analysis of these chemically-induced persister (CIP) cells implicated specific genes in this phenotype, and targeted knockdowns confirmed roles for three genes in either promoting or mitigating the tolerance state. These findings highlight that chemical disruption of the pH gradient is a facile and rapid means to induce drug tolerance, offering a potentially useful tool to probe persister biology in TB and other infectious diseases.

## Introduction

*Mycobacterium tuberculosis* (Mtb), the causative agent of tuberculosis (TB), has been a formidable threat to humanity for millennia, owing to its capacity to evade both immune defenses and therapeutic interventions. With hundreds of millions of people estimated to be infected, and about 1.3 million fatalities in 2022 alone, including nearly one-third of all deaths associated with antibiotic resistance, Mtb has again surpassed COVID-19 as the deadliest infectious pathogen today ^1,2^. Current anti-TB treatments require at least four drugs—rifampicin, isoniazid, ethambutol, and pyrazinamide—often administered over six months in case of drug-susceptible infections. Despite the prolonged and intense nature of these drug regimens, they sometimes fail to eliminate TB, as about 5% of patients suffer treatment failure or relapse ^3-5^. These challenges are further heightened by the toxicities often associated with TB medications and the resulting poor adherence to treatment schedules ^6-8^.

A growing body of evidence attributes TB drug failure and lengthy treatment to a subpopulation of drug-tolerant cells termed “persisters” that withstand bactericidal antibiotic pressure in the absence of known heritable resistance mechanisms ^9-12^. In addition, data from recent studies show an important role of persister cells in the evolution of antibiotic resistance ^13-16^. The concept of antibiotic persistence as a survival strategy distinct from genetic resistance emerged from the pioneering work by Gladys Hobby and Joseph Bigger in 1940s ^10,17^. Hobby first observed this phenomenon in a streptococcal culture, noting that a small percentage of bacteria survived penicillin treatment. Joseph Bigger subsequently built upon this observation, coining the term “persisters” and further documenting their existence in *Staphylococcus aureus* cultures. Both studies showed that persisters typically exist at a very low frequency *in vitro*. Furthermore, Bigger postulated that the ability of a minority of bacterial cells to survive antibiotic killing was not due to heritable resistance but rather a transient tolerance ^17^. Bigger’s insights spurred decades of research aimed at elucidating the underpinnings of this phenomenon. These studies have identified several mechanisms that contribute to persistence, such as metabolic dormancy, low ATP levels, efflux hyperactivity, accumulation of misfolded proteins, and activity of toxin-antitoxin modules ^9, 18-24^. Yet, despite these advances, our understanding of these mechanisms and the interplay between them in triggering and maintaining the persistence state remains far from complete.

To help deconvolute the biology of TB persisters and the underlying mechanisms, we employed a chemical screen to identify small molecules that target persisters. Surprisingly we discovered that the FDA-approved anthelmintic drug niclosamide (NCA) exhibits broad-spectrum drug tolerance- *promoting* activity. Previous studies have reported NCA as a promising anti-TB drug that exhibited activity against Mtb cells through disruption of the proton motive force (PMF) ^25 26, 27^. However, our data reveal unexpected negative impacts of NCA on the bactericidal activities of current anti-TB drugs. Here we demonstrate that NCA’s disruption specifically of the pH gradient portion of the PMF drives Mtb cells into a low-energy, non-replicating and drug-tolerant state. This chemically induced persistence (CIP) provides a potentially valuable model for probing the molecular mechanisms of TB persistence.

## Results

### Identification of niclosamide as a drug persistence-promoting agent

To uncover chemical modulators of Mtb persistence, we screened 2,336 FDA-approved drugs at single dose for their ability to impact the sterilizing activity of isoniazid (INH) and rifampicin (RIF) against Mtb (H37Rv). In this screen, log-phase H37Rv Mtb adjusted to ∼ 1.7 x 10^7^ CFU/well were treated with INH/RIF at 100x MIC each (22.75/0.49 µM) +/- compounds of the drug library (MIC data are provided in Supplementary Table 1). Given the large Mtb inoculum used (∼5.9 × 10^10^CFU total), we performed the screen in the presence of these two frontline drugs to help mitigate the risk of resistance. At defined times, cells were washed and fresh drug-free 7H9/OADC medium was added to allow for growth recovery, which was then assessed by measuring cellular ATP levels after 6 weeks of incubation at 37°C (Fig. 1a). We recently published a 7-day version of this screen that identified compounds with anti-persister activity ^28^. Here we describe results after 14 days of drug treatment, in which INH/RIF alone was sterilizing, so we were unable to identify additional anti-persister molecules with this assay. However, surprisingly, we detected two compounds, niclosamide (NCA) and oxybuprocaine, that interfered with the sterilizing activity of INH/RIF. Unlike the rest of the small molecule library, Mtb cells treated with either compound in combination with INH/RIF restored ATP levels after six weeks in drug-free medium (Fig. 1b). NCA showed a stronger protective phenotype compared to oxybuprocaine and was therefore selected as the primary focus for subsequent study.

**Figure 1.**
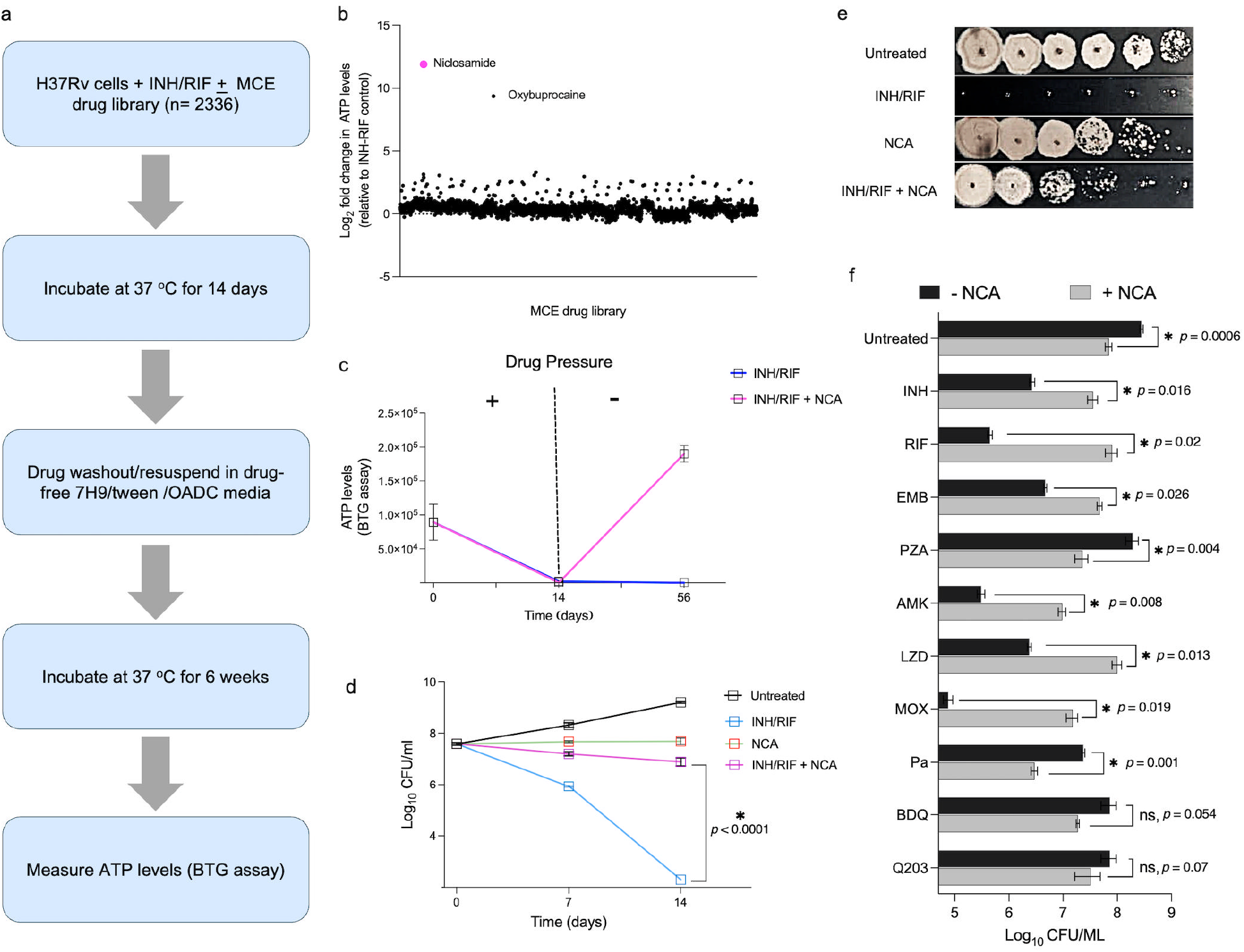
Identification of niclosamide (NCA) as a persistence-promoting agent in *Mycobacterium tuberculosis*(a) Schematic representation of the screening platform as described in the method section. (**b**) Screening results demonstrate that NCA-treated Mtb cells (H37Rv) survive the sterilizing effect of INH/RIF (isoniazid-rifampicin), as evidenced by ATP level rebound six weeks after drug withdrawal. (**c**) Temporal measurements of ATP levels in response to NCA at 2x MIC in the presence or absence of INH/RIF (100x MIC). (**d**) Time-kill analysis depicting CFU in response to the indicated treatments. The asterisk indicates statistical significance, and the *p*-value was determined by two-way ANOVA followed by Dunnett’s multiple comparisons test. (**e**) Spot assays showing CFU growth on 7H10 plates after treating Mtb H37Rv cells with NCA (2x MIC), INH/RIF (100x MIC), or both for 14 days. (**f**) Effect of NCA on the anti-TB activity of INH, RIF, EMB (ethambutol), PZA (pyrazinamide), AMK (amikacin), LZD (linezolid), MOX (moxifloxacin), Pa (pretomanid), BDti (bedaquiline), and ti203 (telacebec). Mtb (H37Rv) cultures were treated for 28 days with the indicated drugs at 10x MIC, except PZA was tested at 500 μM in the presence or absence of NCA (2x MIC). Asterisks indicate statistical significance and *p*-values were determined by a two-tailed Welch’s t-test.

NCA is an FDA-approved antihelmintic previously reported to have promising anti-TB activity ^25-27^. We confirmed that NCA inhibits growth of log-phase Mtb in vitro, with an MIC of 4 µM. (Supplementary Table 1). In our persistence assay, we found that INH/RIF+NCA treatment reduced ATP levels to background by day 14, however, upon removal of drug pressure, cells were able to restore their ATP levels, indicating recovery of cell growth (Fig. 1c). To validate the persistence-promoting activity of NCA, we performed a time-kill study. We found that NCA at 2x MIC exhibited bacteriostatic activity, even in the presence of 100x MIC of INH/RIF; however, INH/RIF alone completely sterilized the culture (Fig. 1d, e). To determine whether NCA’s protective activity is due to reduced TB drug uptake, we measured intracellular RIF levels in the presence and absence of NCA. Mass spectrometry analysis revealed comparable intracellular RIF concentrations between NCA-treated and untreated Mtb cells (Supplementary Figure 3), indicating that NCA does not affect intracellular drug accumulation.

**Figure 2.**
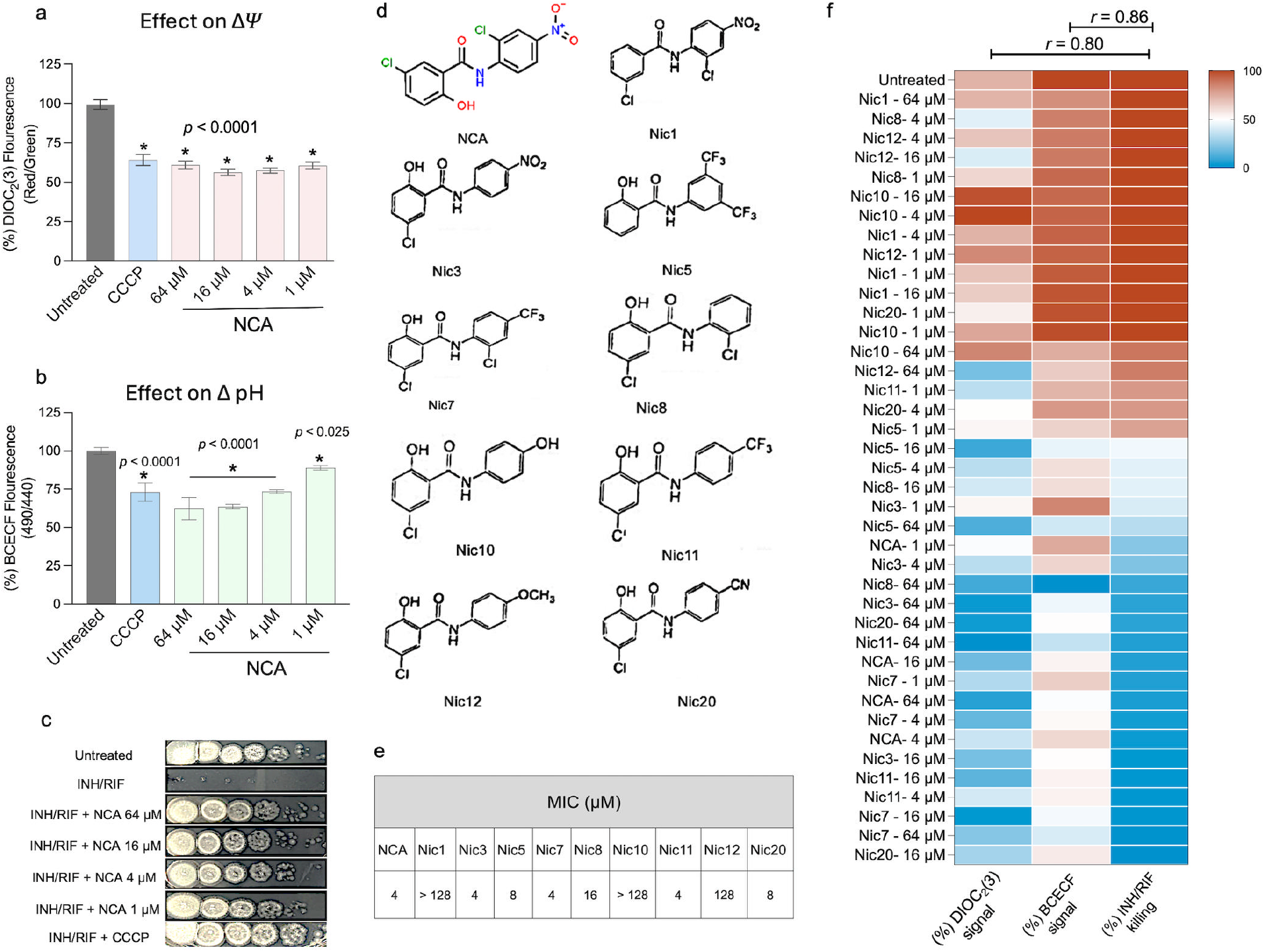
Disruption of the proton motive force (PMF) is correlated with INH/RIF tolerance in *M. tuberculosis*. (**a**) Effect of NCA on the membrane potential, measured by the DIOC_2_(3) assay. (**b**) Effect of NCA on the intracellular pH, measured by the BCECF assay. (**c**) Spot assays showing Mtb (H37Rv) survival after 14-day treatment with INH/RIF (100× MIC) in combination with varying concentrations of NCA or with CCCP (64 μM). (**d**) Chemical structures of NCA analogs. (**e**) MIC data of NCA and its analogs against H37Rv. (**f**) Heatmap illustrating the effect of NCA and its analogs on PMF and INH/RIF killing along with corresponding Pearson correlation coefficients (*r*). Asterisks denote statistical significance; *p*-values were determined by ordinary one-way ANOVA with Dunnett’s post-hoc test for multiple comparisons.

**Figure 3.**
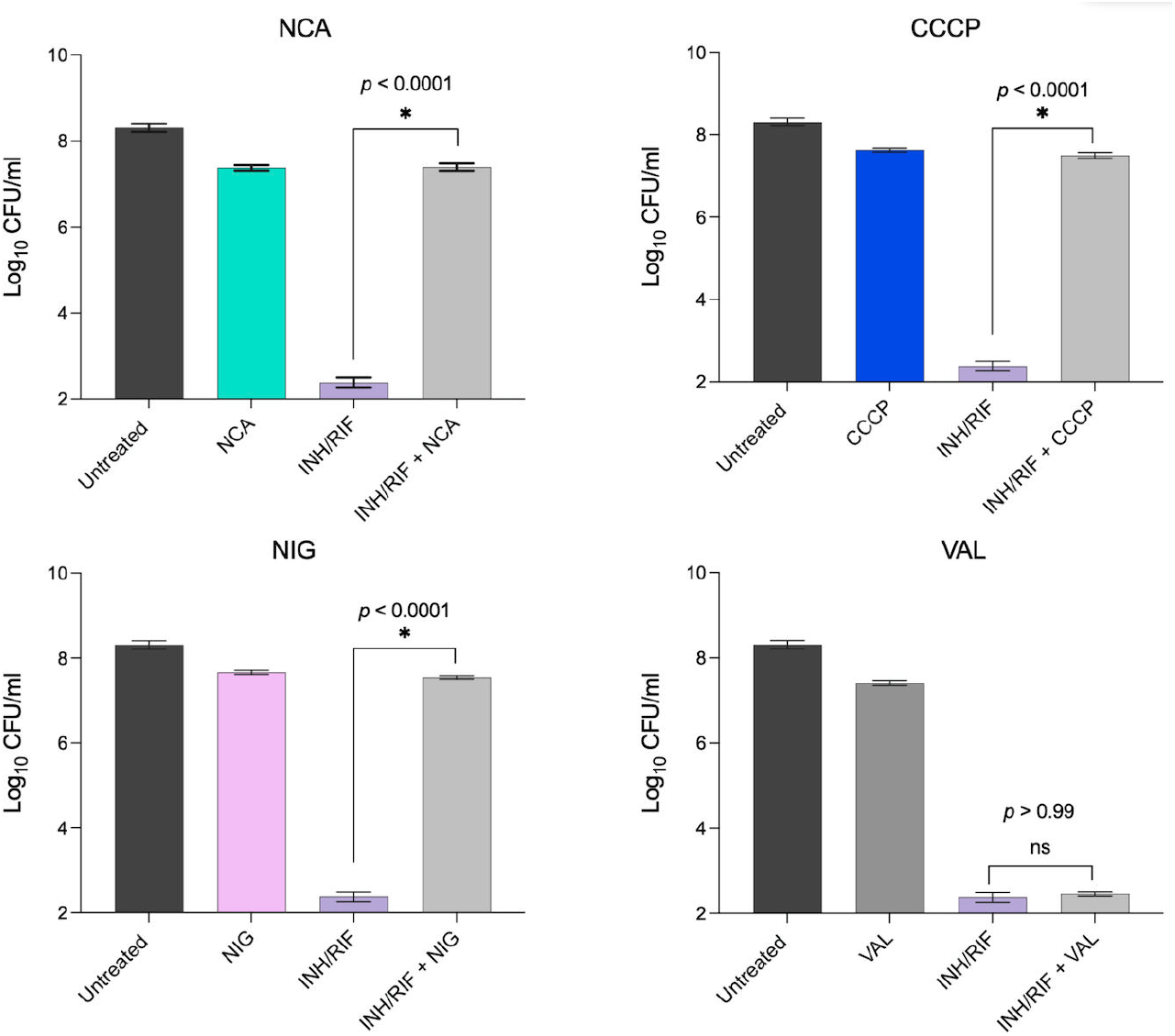
Effect of different PMF inhibitors on INH/RIF killing activity. H37Rv cultures were treated for 14 days with INH/RIF (100x MIC) + 2x MIC of the indicated drugs: niclosamide (NCA), CCCP, nigericin (NIG), and valinomycin (VAL). Data represent mean log10 CFU/ml ± standard deviation from three replicates. Asterisks indicate statistical significance and *p*-values were determined by a two-tailed independent Welch’s t-test.

Next, to investigate whether the protective effect of NCA is TB-drug specific, we examined its interactions with 10 different anti-TB agents. Our findings showed that NCA significantly hindered the bactericidal activity of INH, RIF, ethambutol (EMB), amikacin (AMK), linezolid (LZD), and moxifloxacin (MOX), leading to an increase in the surviving CFU by 1–3 logs (*p* < 0.05) (Fig. 1f). Conversely, NCA enhanced the killing activity of pyrazinamide (PZA) and pretomanid (Pa), resulting in a reduction of surviving CFUs by approximately 1 log. We also observed a modest, non-significant CFU reduction of approximately 0.5 log when NCA was combined with the ATP synthesis inhibitors bedaquiline (BDQ) and telacebec (Q203), relative to single-agent treatments (Fig. 1f). Additionally, we investigated whether NCA could influence the intracellular killing activity of RIF using a THP-1 infection model. Our data showed that NCA treatment at only 0.25x MIC resulted in a substantial interference in RIF’s intracellular killing activity (Supplementary Figure 1). Then, since persistence has been noted in many bacterial pathogens, we tested whether NCA could promote drug tolerance in other species. We found that NCA protects both *E. coli* and *S. aureus* from otherwise lethal doses of antibiotics (Supplementary Figure 2).

### NCA-induced drug tolerance is associated with its ability to disrupt the proton motive force

Prior studies have shown that the anti-TB activity of NCA is due to its capacity to disrupt the two components of proton motive force (PMF), membrane potential (ΔΨ) and the proton gradient (ΔpH) ^25, 26^. Thus, we aimed to investigate whether there is a connection between PMF disruption and the drug-tolerance promoting activity of NCA. We first assessed the effect of varying concentrations of NCA (1–64 μM) on ΔΨ using the membrane-potential-sensitive dye DIOC_2_(3) ^25, 29, 30^, and on the pH gradient using the pH-sensitive dye BCECF ^25, 31, 32^. Across all tested concentrations, NCA induced a significant membrane depolarization, and depletion of the pH gradient, similar to the known PMF inhibitor, CCCP (Fig. 2a and b). To test whether these changes were linked to altered drug tolerance, we measured the ability of NCA at different concentrations to promote survival in the presence of INH/RIF. We found that NCA triggered a strong tolerance phenotype at all tested concentrations, and that CCCP also elicited drug tolerance (Fig. 2c). These findings suggest that the ability of these compounds to promote antibiotic tolerance may be associated with their capacity to disrupt the PMF.

To further assess this correlation, we evaluated a series of nine structurally diverse NCA analogs (Fig. 2d) for their anti-TB activities, PMF modulation, and effects on INH/RIF killing. Using the broth microdilution method, we found that these NCA analogs demonstrated varied antibacterial activities against Mtb. As shown in Fig. 2e, NCA contains two phenyl rings connected by an amide group with different substituents. The hydroxyl group (OH) on the A ring is essential for activity; removing this group, as in Nic1, eliminates activity. On the other hand, the nitro (NO2) and chlorine (Cl) groups on the B ring are not essential and can be substituted with other electron-withdrawing groups, allowing analogs such as Nic3, Nic5, Nic7, Nic8, Nic11, and Nic20 to retain different degrees of inhibitory activity. Compounds Nic3, Nic5, Nic7, Nic11, and Nic20 were found to have comparable activity to NCA with MIC values ranging from 4 to 8 μM, while Nic8 exhibited moderate activity (MIC = 16-32 μM). Substituting the B ring with electron-donating groups led to inactive compounds, as observed in Nic10 and Nic12 (MIC > 128 μM).

Next, we evaluated the impact of NCA analogs on the PMF and INH/RIF killing. Our results demonstrate that the disruption of ΔΨ and ΔpH correlates well with both the antibacterial effects and tolerance to INH/RIF. Analogs with potent anti-Mtb activities exhibited significant disruption of ΔΨ and ΔpH and conferred tolerance to INH/RIF, while NCA analogs with weak antibacterial activity displayed minimal or no effects on the PMF and failed to interfere with INH/RIF killing activity (Fig. 2f). Overall, there was a strong correlation between ΔΨ or ΔpH disruptions and tolerance to INH/RIF (*r* = 0.80, and 0.86 respectively).

### Disruption of the proton gradient is key to NCA’s protective activity

While these findings suggest a link between PMF disruption and drug tolerance, whether a specific PMF component was driving this effect remained unclear. To investigate this question, we compared the protective effects of NCA and CCCP with those of nigericin and valinomycin—two specific PMF inhibitors with distinct mechanisms of action. Nigericin is a disruptor of the proton gradient by facilitating K^+^ /H^+^ exchange across the membrane, leading to intracellular acidification without compromising the membrane potential ^3334^. On the other hand, valinomycin specifically facilitates K^+^ transport, resulting in depolarization of the cell membrane without reducing the proton gradient ^35,36^ . We tested the ability of both compounds to influence drug susceptibility in Mtb, and found that nigericin, like NCA and CCCP, conferred a significant tolerance to INH/RIF, while conversely valinomycin did not (Fig. 3). These results indicate that disruption of ΔpH, rather than ΔΨ, is the key PMF component involved in the observed drug tolerance phenotype.

### pH-dependent modulation of NCA-mediated drug tolerance

Our findings indicate that intracellular pH is a key determinant of the Mtb response to antibiotic pressure. We therefore hypothesized that manipulation of extracellular pH in the presence of NCA would differentially alter Mtb susceptibility to INH/RIF. To test this, Mtb cultures were incubated under three defined pH conditions—acidic (pH 6.0), neutral (pH 7.0), and alkaline (pH 8.0)—and exposed to INH/RIF (100×MIC) for 7 days, with or without NCA at either a subinhibitory concentration (0.5×MIC) or a suprainhibitory concentration (10×MIC).

At the low NCA dose, maximal protection was observed under acidic conditions (pH 6.0), with a 1.53 log increase in CFU compared to the INH/RIF control (Fig. 4a). Protection was modest at neutral pH (0.7 log increase) and was completely lost at alkaline pH (pH 8.0), consistent with the requirement for proton influx and consequent cytosolic acidification in mediating the NCA protective effect. In contrast, at the high NCA dose, protection was greatest at neutral pH (1.7 log increase), absent at pH 8.0, and notably reversed to enhanced killing at pH 6.0, where CFU counts were reduced by approximately 1 log relative to the INH/RIF group, (Fig. 4b). These data suggest that at alkaline pH, reduced proton availability limits the capacity of NCA to acidify the cytosol, thereby preventing induction of the drug-tolerant state. Conversely, under acidic conditions, high-dose NCA may cause excessive cytosolic acidification that surpasses the bacterium’s tolerance threshold, resulting in cell death. Collectively, these results reveal a biphasic relationship between intracellular acidification and antibiotic tolerance, in which protection is achieved only within a defined range of pH perturbation. Outside this range, tolerance is lost and, in extreme acidification, replaced by bactericidal activity.

**Figure 4.**
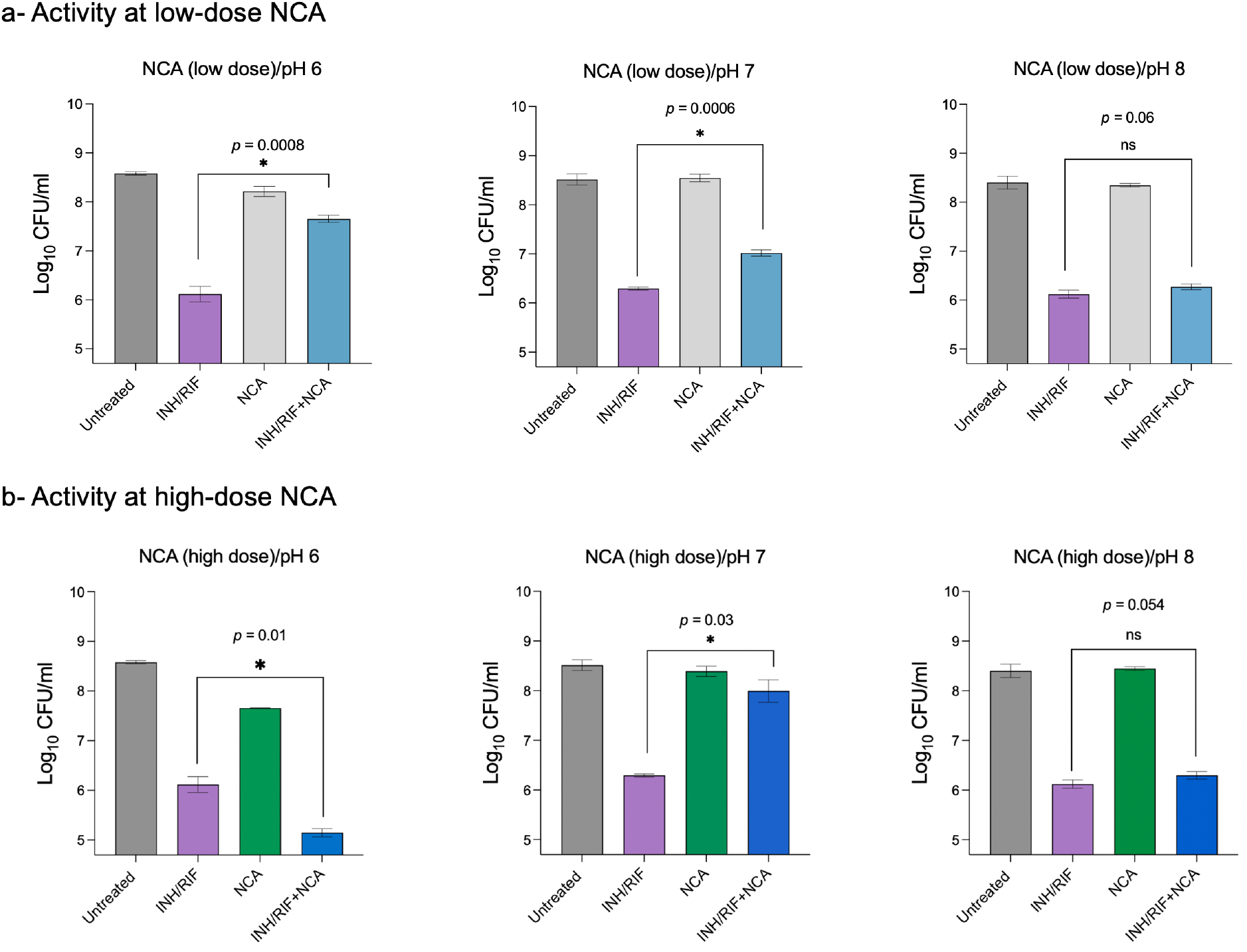
External pH modulates the protective activity of NCA against INH/RIF. (a)Effect of low-dose NCA (0.5×MIC) on Mtb H37Rv survival in the presence or absence of INH/RIF (100×MIC) under acidic (pH 6.0), neutral (pH 7.0), and alkaline (pH 8.0) conditions. (b)Effect of high-dose NCA (10×MIC) under the same treatment and pH conditions. Data represent mean log10± standard deviation from three biological replicates. Asterisks denote statistical significance and *p*-values were determined using a two-tailed independent *t*-test.

### NCA, CCCP, and nigericin elicit similar transcriptomic changes compared to valinomycin

To gain a deeper understanding of how PMF inhibition contributes to INH/RIF tolerance, we performed RNA-seq to examine the transcriptomic changes in response to treatments with NCA, CCCP, nigericin, and valinomycin. Even at 0.5x MIC for only one hour, these PMF inhibitors exerted a significant impact on the Mtb transcriptome. The number of differentially expressed genes showing significant (*p* < 0.05) upregulation relative to the untreated control ranged from 289 to 1,043, while significantly downregulated genes ranged from 500 to 1,696, underscoring the substantial stress induced by these inhibitors (Fig. 5a). Additionally, comparing the transcriptomic profiles of Mtb treated with NCA, CCCP, and nigericin to each other revealed strong positive correlations, with Pearson correlation coefficients (*r*) ranging from 0.74 to 0.92 (Fig. 5b). However, these correlations dropped to below 0.5 when we compared these profiles to that of Mtb treated with valinomycin. These findings support the idea that NCA, CCCP, and nigericin may induce tolerance via a shared mechanism.

**Figure 5.**
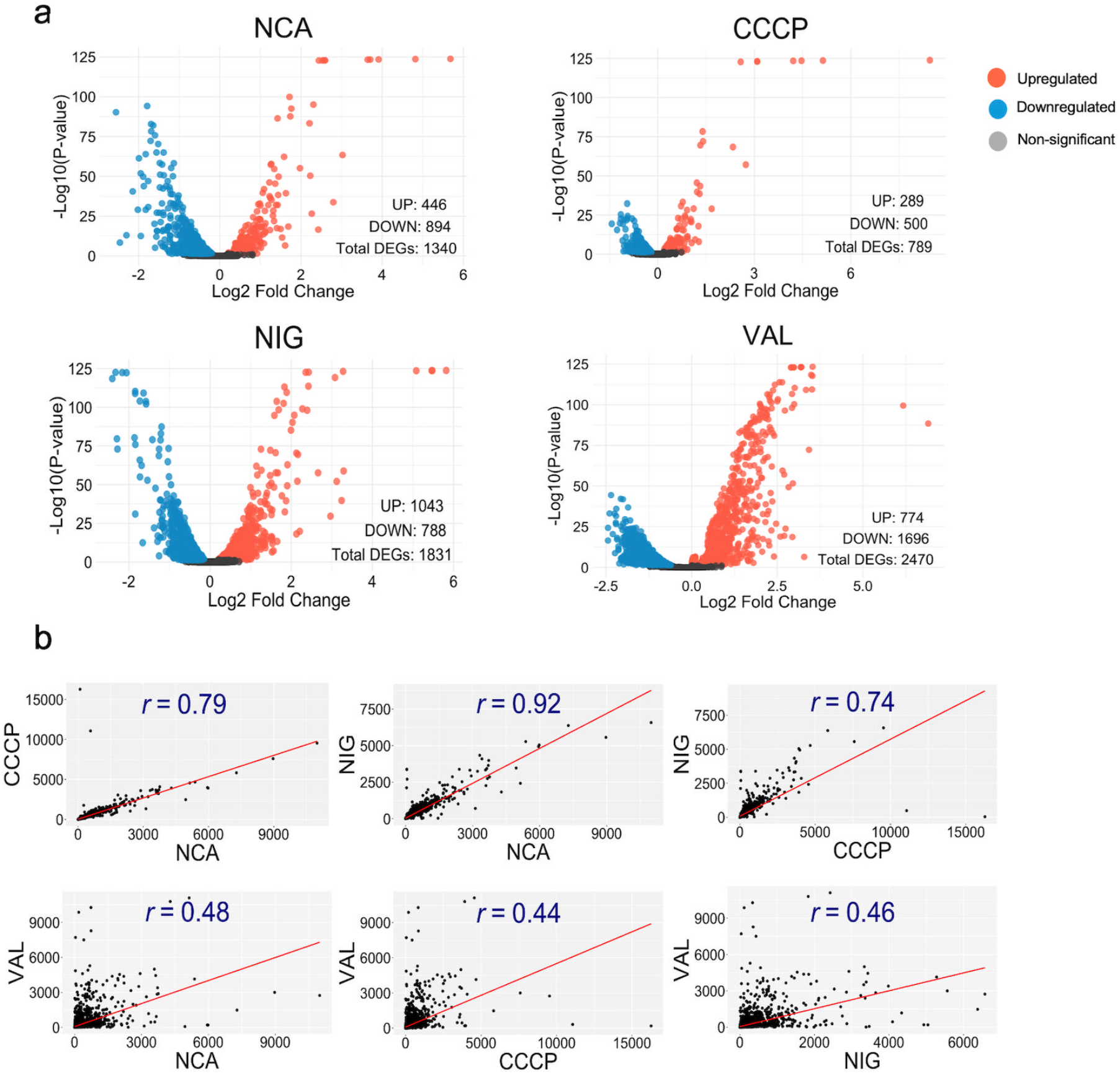
Transcriptomic profiling of PMF inhibitors in *Mycobacterium tuberculosis*. (**a**) Volcano plots illustrating the effect of the indicated PMF inhibitors (at 0.5x MIC) on Mtb (H37Rv) transcriptome. The plots highlight the number of significantly differentially expressed genes (*p* < 0.05) compared to the untreated control, (**b**) Scatter plots showing Pearson correlation coefficients calculated for the transcriptomic profiles of tested PMF inhibitors. Transcripts per million (TPM) were used to generate the scatter plots.

### Identification of key genes and significantly enriched pathways in PMF-disrupted drug-tolerant Mtb cells

Our phenotypic assays and transcriptomic data suggest that, unlike valinomycin, NCA, CCCP, and nigericin may converge on a common pathway that promotes tolerance to anti-TB drugs, despite differences in their structures and primary modes of action. To dissect this mechanism, we focused on genes and pathways commonly regulated only by the tolerance-promoting agents but not by valinomycin. Sixteen transcripts significantly regulated (≥ 1 log_2_ fold change vs. untreated control; *p* < 0.05) by NCA, CCCP, and nigericin, but not by valinomycin, were identified — 10 upregulated and 6 downregulated (Fig. 6a, Supplementary Table 2). Of these, 11 transcripts showed at least a 2-fold difference relative to valinomycin (Fig. 6b). Our results show that Rv3290c and Rv3289c were the most strongly upregulated genes in response to tolerance-promoting inhibitors, followed by Rv1057, Rv2428, and Rv2107, compared to valinomycin. In contrast, Rv2987c, Rv2988c, Rv0057, Rv0059, Rv0241c, and Rv0242c were downregulated by the tolerance-promoting inhibitors but upregulated by valinomycin (Fig. 6c).

**Figure 6.**
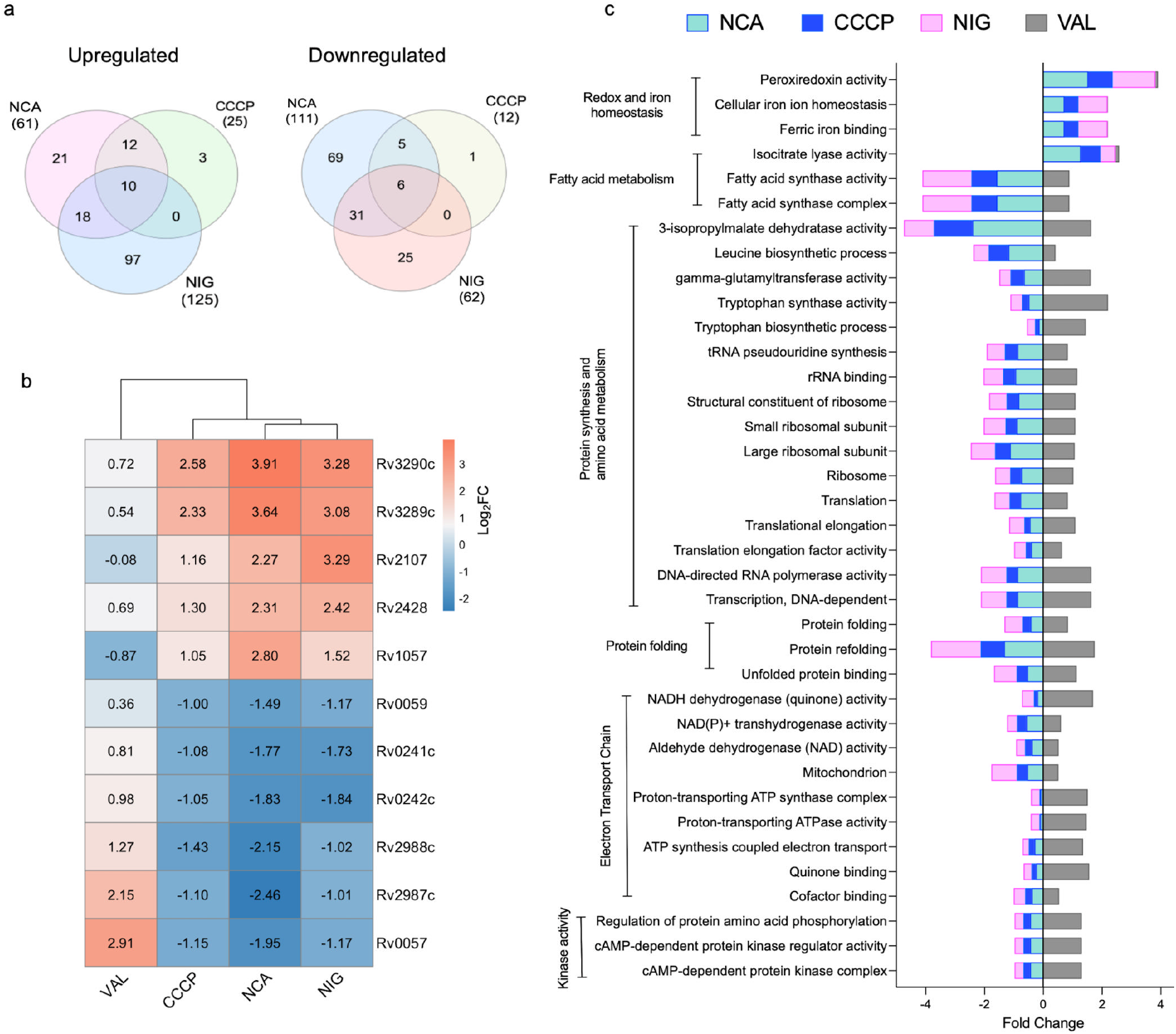
Shared genes and gene sets among tolerance-promoting PMF inhibitors (**a**) Venn diagrams showing the number of DEGs (p < 0.05, ≥1 log_2_ fold change) that were up-or downregulated in the NCA, CCCP, and NIG (nigericin) groups relative to the untreated control. (**b**) Heatmap showing log_2_ fold changes of genes co-regulated by NCA, CCCP, and NIG, with ≥1 log_2_ difference from VAL. VAL values included for comparison. (**c**) Gene ontology analysis showing GO terms representing biological processes that are significantly over/underrepresented (*p* < 0.05) in response to the treatment with the tolerance-promoting PMF inhibitors.

We performed gene enrichment analysis to identify biological processes and pathways significantly affected in PMF-disrupted, drug-tolerant Mtb cells. Pathways related to oxidative stress response, iron ion homeostasis, and isocitrate lyase activity were significantly enriched (p < 0.05) in cells treated with NCA, CCCP, and nigericin, but not with valinomycin, compared to the untreated control. Conversely, energy-demanding pathways such as amino acid biosynthesis, protein translation, and fatty acid synthesis were significantly downregulated only in the presence of the tolerance-promoting PMF inhibitors (Fig. 6c).

### Genes with a role in Mtb persistence

To investigate the potential role of selected differentially expressed genes in Mtb drug tolerance, we utilized targeted CRISPR interference (CRISPRi) to knock down the 11 genes, identified as being co-regulated by drug tolerance-promoting inhibitors but not by valinomycin. These conditional knockdown mutants were induced with anhydrotetracycline (ATC) and their response to INH/RIF treatment was evaluated. Our analysis identified three knockdown strains with altered responses to INH/RIF treatment. Knockdown of Rv0057, a gene repressed under tolerance-promoting conditions, led to enhanced Mtb survival in the presence of INH/RIF compared to the uninduced control (Fig. 7a). In contrast, knockdown of Rv2428 and Rv3289c, both upregulated under tolerance-promoting conditions, resulted in increased susceptibility to INH/RIF (Fig. 7a). To assess the contribution of these genes to the cellular bioenergetics, we measured intracellular pH and ATP levels. We found that only Rv0057 knockdown was associated with a significant reduction in intracellular pH, exhibiting a comparable effect to the positive control (nigericin/K^+^/ pH 5.5) (Fig. 7b). In contrast, knockdown of Rv2428 nor Rv3289c did not significantly affect intracellular pH (Fig. 7b). Additionally, we found that both Rv0057 and Rv3289c knockdown strains showed significantly lower ATP levels compared to the wild-type H37Rv strain (Fig. 7c).

**Figure 7.**
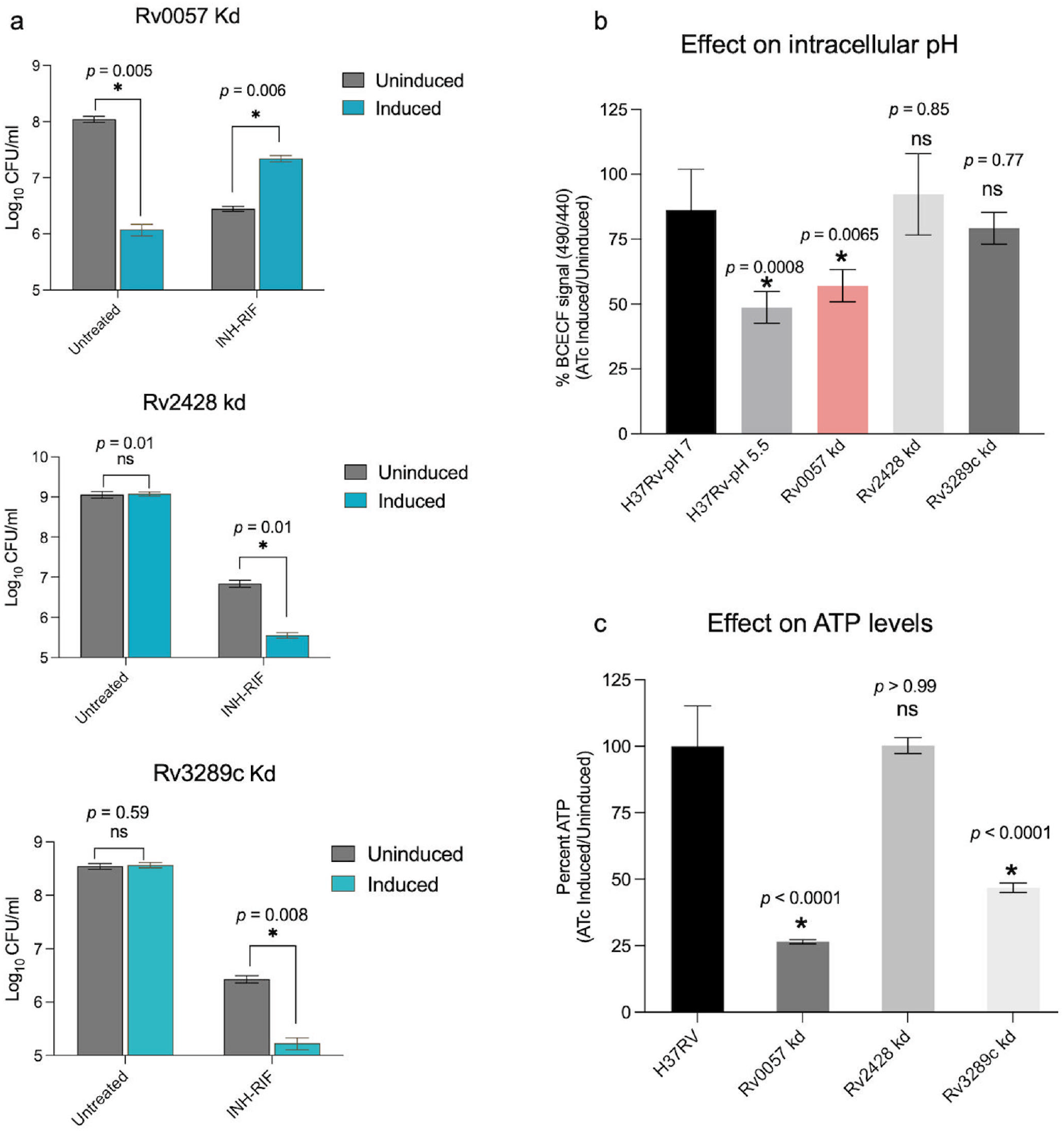
Genes with a role in persistence. (a) Effect of 7-day treatment with INH/RIF (100x MIC) on Rv0057, Rv2428, and Rv3289c knockdown strains. Asterisks indicate statistical significance and *p*-values were determined by a two-tailed independent Welch’s t-test. Measurements of (b) intracellular pH using the BCECF assay and cellular ATP levels (c) in Rv0057, Rv2428, and Rv3289c knockdown strains. Asterisks denote statistical significance; *p*-values were determined by ordinary one-way ANOVA with Dunnett’s post-hoc test for multiple comparisons.

## Discussion

Mtb persisters present a substantial challenge to the effective control of TB, contributing to infection relapse, prolonged treatment time, and the emergence of drug-resistant isolates ^37-39^. The scarcity and transitory nature of persister cells, along with non-genetic variability and the dynamic equilibrium between persistent and non-persistent states, have made it challenging to dissect the underlying mechanisms of this phenomenon ^40-42^. Here, we characterize an unexpected effect of the FDA-approved anthelmintic drug niclosamide (NCA) on Mtb persistence. Despite previous reports of potent anti-Mtb activity, we found that NCA paradoxically rendered H37Rv cells tolerant to a broad range of anti-TB drugs, including isoniazid, rifampicin, ethambutol, amikacin, linezolid, and moxifloxacin. We term this phenomenon *chemically induced persistence* (CIP), as NCA treatment drove Mtb into a low-ATP, non-replicating, drug-tolerant state that mirrors hallmark features of persister cells. Upon recovery, NCA-protected cells displayed unchanged drug susceptibility profiles (data not shown), consistent with a non-genetic basis for this phenotype. The CIP model provides a practical tool to study persister biology, as NCA treatment effectively enriches for a drug-tolerant population, rendering nearly the entire culture tolerant. Further, we found that NCA treatment elicits drug tolerance in *E. coli* and *S. aureus*, indicating that its mechanism is broadly conserved.

We hypothesized that NCA’s protective activity arises from its ability to disrupt proton motive force (PMF). This hypothesis was supported by testing a panel of NCA analogs for their effects on PMF and their influence on INH/RIF susceptibility. Our data revealed a strong correlation between disruption of ΔΨ and ΔpH and the level of drug tolerance (*r* = 0.8 and 0.86, respectively). To further validate this mechanism, we evaluated other known PMF disruptors, including CCCP, nigericin, and valinomycin. Like NCA, both CCCP and nigericin—but not valinomycin—induced a comparable drug tolerance phenotype. Since nigericin selectively disrupts ΔpH and valinomycin specifically interferes with ΔΨ, this finding indicates that disruption of ΔpH, rather than ΔΨ alone, plays a key role in driving this tolerance phenotype. In addition, we found that NCA’s activity was tuned by extracellular pH: at subinhibitory concentrations, protection was maximal under acidic conditions, modest at neutral pH, and lost at alkaline pH, consistent with a requirement for proton influx. At higher concentrations, excessive acidification under acidic conditions became bactericidal, revealing a biphasic relationship in which tolerance occurs only within a defined range of intracellular pH. Together, these findings show that ΔpH disruption, modulated by extracellular pH, is central to NCA-mediated drug tolerance.

Supporting evidence for the role of intracellular acidification in this process can also be found in other bacterial systems. For example, Bartek et al. ^43^ showed in *M. smegmatis* that conditions favoring proton entry were associated with reduced antibiotic lethality. Similarly, studies in *E. coli* have demonstrated that persister cells, comprising ∼1% of stationary phase populations, maintain a more acidic intracellular environment compared to drug-susceptible wild-type cells ^44^. Furthermore, Thompson et al. ^45^ recently reported that the anti-tolerance effect of adenosine in *E. coli* was linked to its ability to modulate PMF and induce intracellular alkalinization. Together, our findings—along with prior reports from diverse bacterial species—suggest that collapse of the pH gradient, despite its fitness cost, may serve as a survival strategy under stress. These insights may have clinical implications, as *M. tuberculosis* is known to inhabit acidic environments within host macrophages and granulomas. Under such conditions, TB bacilli experience acid stress that may disrupt their pH gradient, potentially enhancing drug tolerance *in vivo*.

To gain deeper insights into the biology of PMF-disrupted persister cells, we performed RNA sequencing to identify key genes and transcriptional adaptations associated with drug tolerance. This analysis revealed upregulation of genes involved in peroxidase and peroxiredoxin functions, indicating that enhanced oxidative stress responses may be a hallmark of these persister cells— consistent with prior reports linking redox defense mechanisms to bacterial persistence ^46-48^. We also observed upregulation of genes associated with isocitrate lyase activity, an enzyme previously shown to divert acetyl-CoA from fatty acid β-oxidation into the glyoxylate shunt. This metabolic shift allows for net carbon gain and may support the survival of persister cells *in vivo* by providing an alternative energy source ^49^. In contrast, genes involved in electron transport chain activity and several energy-demanding processes—such as protein synthesis, fatty acid biosynthesis, and protein folding—were significantly downregulated, suggesting that bioenergetic output is tightly regulated under drug-tolerant conditions. At the gene level, we identified 11 genes that were similarly regulated following treatment with NCA, nigericin, and CCCP, but not valinomycin. To evaluate the functional contribution of these genes to INH/RIF tolerance, we used targeted CRISPRi knockdown. This analysis identified three genes with roles in the persister phenotype: *Rv2428* (*ahpC*), *Rv3289c*, and *Rv0057. AhpC*, encoding alkyl hydroperoxidase, is a well-characterized component of Mtb’s antioxidant defense system and has been implicated in INH resistance ^50-53^ . Mutations in the *oxyR-ahpC* intergenic region are known to compensate for *katG* loss-of-function mutations in resistant strains ^54^. In our study, knocking down *ahpC* significantly reduced tolerance to INH/RIF, suggesting a role for oxidative stress management in Mtb drug response. *Rv3289c*, a putative transmembrane protein of unknown function within the *lat* operon, also emerged as a mediator of drug tolerance. Previous studies reported a robust upregulation of *lat* operon under starvation conditions ^55^, supporting a possible role in persistence. Knockdown of *Rv3289c* both enhanced susceptibility to INH/RIF and decreased intracellular ATP. Reduced ATP has previously been linked to persistence ^56-58^, yet our data show that this correlation does not always hold. Similarly, we recently reported that the antiemetic agent netupitant, which selectively kills persister cells, lowers ATP without affecting ΔpH ^28^. These data support a model in which a dissipated proton gradient and the resulting intracellular acidification, rather than ATP depletion alone, are key drivers of persistence. In contrast, knockdown of *Rv0057*, a hypothetical protein, resulted in increased Mtb survival under INH/RIF treatment. This phenotype was associated with enhanced intracellular acidification and reduced ATP levels—resembling the biological state observed following NCA treatment—suggesting that repression of *Rv0057* may contribute to drug tolerance through bioenergetic remodeling. Together, these findings support a model in which ΔpH disruption triggers coordinated transcriptional and metabolic responses that promote persistence. Further studies are needed to see whether the role of these genes in persistence is drug-specific. However, targeting redox balance and bioenergetics may offer new avenues to enhance treatment efficacy and prevent relapse.

In conclusion, this study reveals that disruption of the proton gradient strongly influences Mtb drug tolerance. While it broadly promotes tolerance to most anti-TB drugs, it enhances susceptibility to PZA and pretomanid—two drugs shown to reduce relapse rates clinically ^59, 60^. This suggests that our CIP model provides a physiologically relevant tool for probing persister biology and screening new therapeutics. Moreover, because fundamental energy-generating processes are targeted, our findings have broad implications for the study of drug tolerance in other bacterial pathogens where antimicrobial resistance remains a major challenge. Future studies are needed to elucidate how intracellular acidification drives drug tolerance at the molecular level, whether it is sufficient to induce persistence on its own, and to characterize ΔpH-disrupted persisters in their native host environments.

## MATERIALS and METHODS

### Bacterial Strains, Culture Conditions, and Reagents

*Mycobacterium tuberculosis* H37Rv was the primary strain used in study and was used as the parental strain for all CRISPRi knockdown mutants. Erdman and HN878 strains were provided by Dr. Rhea Coler’s lab. Mtb Cultures were grown in Middlebrook 7H9 broth supplemented with 10% OADC, 0.2% glycerol, and 0.05% Tween 80 (referred to as 7H9-rich medium), or on 7H10 agar with 10% OADC and 0.2% glycerol. Cultures were incubated at 37 °C on a rotary shaker or wheel incubator. All drug treatments were performed on static cultures. Strains were stored in 15% glycerol at −80 °C. For selection, kanamycin was used at 25 µg/mL. CRISPRi induction was performed using 100 ng/mL anhydrotetracycline (ATc) added 24 h prior to drug treatment. pLJR96 was obtained from Addgene (MA, USA). BsmBI and T4 DNA ligase were purchased from New England Biolab (MA, USA). Rifampicin, isoniazid, ethambutol, moxifloxacin, and amikacin were obtained from Sigma-Aldrich (MO, USA) or Fisher Scientific (NH, USA). Compounds from the MCE FDA-approved drug library—including niclosamide, CCCP, nigericin, valinomycin, pyrazinamide, linezolid, pretomanid, Q203, and bedaquiline—were purchased from Med Chem Express (NJ, USA). ATc was from Acros Organics (MA, USA); kanamycin A from GoldBio (MI, USA); and niclosamide analogs were provided by Dr. Pu Kai Li’s lab. Stock solutions were prepared in DMSO and stored at 4 °C, except amikacin and isoniazid, which were dissolved in water.

### Minimum inhibitory concentration (MIC) determinations

MIC determinations were performed according to the CLSI protocol M24Ed3 with minor modifications ^61, 62^. Test agents were 2-fold serially diluted in Middlebrook 7H9-rich medium in a volume of 100 μl in 96-well culture plates. Then, aliquots of 100 μl of ∼ 5x 10^6^ CFU/mL H37Rv strain were added to each well of the drug dilution plates, yielding a final volume of 200 μl. Plates were then incubated at 37°C for 7 days before growth was detected visually and spectrophotometrically at OD_600_. MIC was defined as the minimum concentration that inhibited the growth of Mtb culture by at least 90% relative to the untreated control.

### Drug screen

*M. tuberculosis* H37Rv strain was grown in 7H9-rich medium to OD_600_ ∼0.5. Assay plates were prepared by dispensing 75 μL of 7H9-rich medium into each well of sterile, clear bottom 96-well V-bottom plates (Greiner), supplemented with 40 μM of compounds from the drug library (MCE). All compounds were screened in combination with a final concentration of INH/RIF (22.75/0.49 µM). These concentrations represent 100x MIC of both compounds in standard MIC assays. INH/RIF treatment was used as the negative control, while INH/RIF combined with thioridazine at 100 µg/mL served as the positive control. Equal volumes of the H37Rv culture were inoculated into each well. Plates were sealed and incubated at 37 °C for 14 days. After incubation, plates were centrifuged, supernatants discarded, and pellets washed once with PBS. Fresh drug-free 7H9-rich medium (150 µL/well) was added. Plates were then sealed in double zip-lock bags and incubated at 37 °C for an additional 6 weeks. Cell viability was assessed by measuring total ATP levels using the BacTiter-Glo™ assay (Promega) ^63^.

### Time-kill assay

Time-kill assays were performed as previously described ^64, 65^. Mtb cultures were grown in 7H9-rich medium to mid-log phase (OD_600_ ∼0.25). For CRISPRi strains, kanamycin (25 µg/mL) was added to maintain plasmid selection, and cultures were diluted in fresh medium with or without anhydrotetracycline (ATc, 100 ng/mL) to induce knockdown. After 24 h of induction, test compounds or vehicle controls were added at the indicated concentrations. At the designated time points, samples were collected, serially diluted, and plated on 7H10 agar for CFU enumeration. Colonies were counted after 3 weeks of incubation at 37 °C. In some experiments, spot assays were performed at the final time point by plating 5 µL from 10-fold serial dilutions onto 7H10 agar, followed by 21 days of incubation before imaging.

### Intracellular Killing Assay in THP-1 Macrophages

The impact of NCA on the intracellular activity of rifampicin (RIF) was assessed using a THP-1 macrophage infection model adapted from previously established protocols ^66^. In brief, THP-1 monocytes were maintained in RPMI 1640 medium supplemented with 10% heat-inactivated FBS and 1% L-glutamine. To induce macrophage differentiation, cells were treated with 50 ng/mL PMA for 48 hours at 37°C. Differentiated cells (5 × 10^5^ per well) were infected with *M. tuberculosis* (H37Rv) at an MOI of 1 and incubated for 2–3 hours to allow phagocytosis. After infection, cells were washed with fresh medium to remove non-internalized bacteria. Infected macrophages were then exposed to RIF (1 µg/mL), either alone or in combination with NCA (0.25 µg/mL; 0.25× MIC), and incubated for 4 days at 37°C. At the end of treatment, cells were lysed using 0.1% Triton X-100, and intracellular bacterial burden was quantified by plating serial dilutions of lysates on 7H10 agar supplemented with OADC. CFUs were counted after incubation, and percent survival was calculated relative to the CFU counts at the start of drug exposure.

### Membrane Potential Assay

Membrane potential was measured as previously described ^25, 29, 30^. Mtb (H37Rv) cultures were grown to mid-log phase (OD_600_ ∼0.5), harvested by centrifugation, and resuspended in 7H9 medium (supplemented with 0.2% glycerol, 0.2% dextrose, 0.085% NaCl, and 0.02% Tyloxapol) to OD_600_ ∼1.0. Compounds (NCA or its analogs) were added at final concentrations of 1, 4, 16, or 64 μM. CCCP (64 μM) was used as a positive control for depolarization, and DMSO served as the negative control. Cultures were immediately incubated with 15 μM DiOC_2_ (3) for 20 min at room temperature, washed to remove excess dye, and resuspended in fresh 7H9 medium. Fluorescence was measured using black, clear-bottom 96-well plates on a SpectraMax M5 plate reader (BMG LABTECH), with excitation at 488 nm and emission at 530 nm (green) and 610 nm (red). Membrane potential was expressed as the red-to-green fluorescence ratio. Each condition was tested in triplicate, and experiments were independently repeated at least twice.

### Intracellular pH assay using the BCECF-AM dye

Intracellular pH was measured using the pH-sensitive fluorescent dye BCECF-AM, as previously described ^31,32^. BCECF-AM is converted to its active, pH-sensitive form by intracellular non-specific esterases. Mtb (H37Rv) cultures were pre-treated with the indicated concentrations of all respective inhibitors used in this study and incubated at 37 °C for 30 min. As a positive control, cells were exposed to 7H9 medium buffered at pH 5.5 containing 10 µg/mL nigericin and 25 mM KCl; DMSO was used as a negative control. After pretreatment, 20 µM BCECF-AM was added, and cultures were incubated for an additional 90 min at 37 °C. Fluorescence was measured using dual excitation at 490 nm and 440 nm, with emission recorded at 530 nm. Intracellular pH was expressed as the fluorescence intensity ratio (490 nm/440 nm). Each condition was tested in triplicate, and all experiments were independently repeated at least twice.

### RNA sequencing and analysis

Actively replicating Mtb (H37Rv) cultures (OD_600_ ∼0.15 in 7H9-rich medium) were treated with various PMF inhibitors at 0.5× MIC for 1 hour at 37 °C. Following treatment, cultures were centrifuged, and cell pellets were collected. RNA was isolated as previously described ^61,67,68^. Briefly, cell pellets in Trizol were homogenized using Lysing Matrix B and a FastPrep 120 homogenizer, followed by centrifugation. The supernatant was extracted with chloroform and RNA was precipitated with isopropanol and high salt solution. Purification was done using the RNeasy kit with DNase treatment (Qiagen), and RNA yield was quantified using a Nanodrop. Ribosomal RNA was depleted using the RiboZero kit (Illumina), and mRNA libraries were prepared with the NEBNext Ultra RNA Library Prep Kit (New England Biolabs) and barcoded using NEBNext Multiplex Oligos. Libraries were sequenced on an Illumina NextSeq 500, generating ∼75 million reads per library. Read alignment was done using a custom pipeline with Bowtie 2 ^69,70^. Raw sequencing data have been deposited in the NCBI Sequence Read Archive (SRA) under accession number PRJNA1226648.

### CRISPRi knockdown strains

The CRISPRi strains were constructed according to the method outlined previously ^71^. Briefly, we used the pLJR965 plasmid encoding a tetracycline-inducible dCas9, a tetracycline-inducible specific sgRNA, and a kanamycin-selectable marker. We made the specific sgRNAs by annealing two complementary oligonucleotides targeting the non-template strand of the ORF 3’ of selected PAM (protospacer adjacent motif) sequences. Forward and reverse primer sequences are available in (Supplementary Table 3). pLJR965 was digested with BsmBI and specific sgRNAs were ligated into digested pLJR965 using T4 DNA ligase. The ligation reactions were transformed into competent *Escherichia coli* and sgRNA insertions were confirmed by Sanger sequencing before the plasmids were transformed into Mtb (H37Rv).

### Statistics and Reproducibility

Unless otherwise indicated, experiments were independently repeated at least three times, and data are presented as mean ± standard deviation from biological replicates. Statistical analyses were performed using two-tailed independent t-test, or one-or two-way ANOVA with Dunnett’s post hoc correction for multiple comparisons, as appropriate. Pearson correlation was used to assess linear associations. Statistical significance was defined as *p* < 0.05.

## Code Availability

All R scripts used for data analysis and figure generation, including volcano plots, gene filtering, Venn diagrams, heatmaps, and correlation analyses, are publicly available in the GitHub repository: https://github.com/HassanEldesouky13/PMF-inhibitors-TB-analysis This repository contains all necessary scripts along with instructions for reproducing the results reported in this study.

## Author Contributions

H.E.E. and D.R.S. conceived and designed the project and wrote the manuscript. H.E.E. performed the screening study and conducted most experiments. K.N.A. conducted the THP-1 cell experiments. J.K.B. assisted with sequencing library preparation. L.K.T.A. and M.G. performed experiments involving *E. coli* and *S. aureus*. E.X. and P.-K.L. synthesized the niclosamide analogs and conducted the SAR analysis. All authors reviewed and approved the final manuscript.

## ACKNOWLEDGMENTS

We thank Natalie Gleason for helping with the drug screen, and all members of the Sherman Lab for helpful discussions. We thank the Mass Spectrometry Center at the University of Washington School of Pharmacy for performing the mass spectrometry analysis, and the Rhea Coler Lab for providing the *M. tuberculosis* Erdman and HN878 strains. This work was supported by a sub-award from NIH P30AI168034 to HE, and by NIH U19 AI 1625598 and R01 AI 146194 to DRS.

