## Supplementary Table 1, Supplementary Figure 1, Supplementary Figure 2, Supplementary Figure 3, Supplementary Table 2, Supplementary Table 3 for "The pH gradient contributes to persistence in *Mycobacterium tuberculosis*"

**Supplementary Table 1**: Minimum inhibitory concentrations of different inhibitors against Mtb H37Rv strain.

| MIC (μM) | |
|---|---|
| Isoniazid (INH) | 0.25 |
| Rifampicin (RIF) | 0.005 |
| Ethambutol (EMB) | 0.25 |
| Pyrazinamide (PZA) | > 1000 |
| Amikacin (AMK) | 1 |
| Linezolid (LZD) | 3 |
| Moxifloxacin (MOX) | 0.15 |
| Pretomanid (Pa) | 0.2 |
| Bedaquiline (BDQ) | 0.5 |
| Telacebec (Q203) | 0.004 |
| Niclosamide (NCA) | 4 |
| Carbonyl cyanide 3 chlorophenylhydrazone (CCCP) | 8 |
| Nigericin (NIG) | 8 |
| Valinomycin (VAL) | 1 |

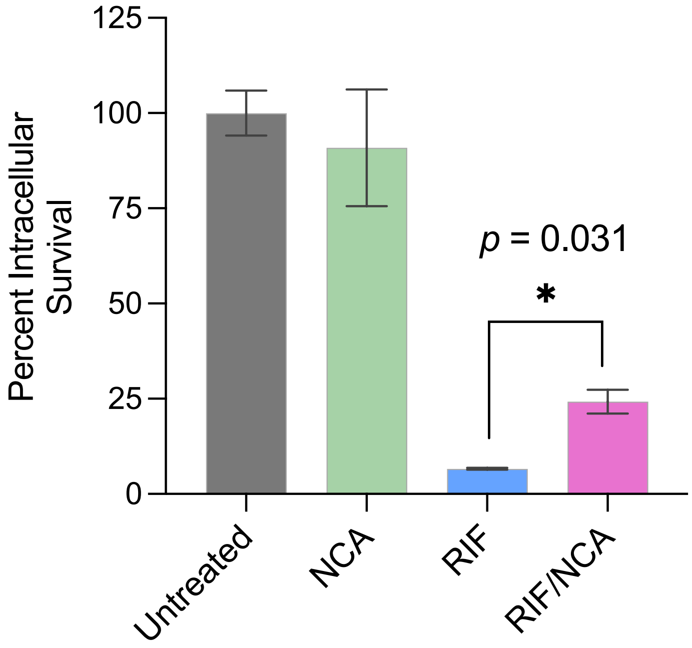


**Supplementary Figure 1. Effect of niclosamide (NCA) on rifampicin (RIF)-mediated intracellular killing of Mtb.** THP-1 cells were infected with *M. tuberculosis* H37Rv at a multiplicity of infection (MOI) of 1:1. Infected cells were then treated with RIF (1 μg/ml), NCA (0.25× MIC), or the combination of RIF and NCA. Data represent the mean percent survival of intracellular Mtb at day 4 post-infection ± standard deviation. Asterisks denote statistical significance; *p*-values were determined using a two-tailed Welch's *t*-test (n=4).

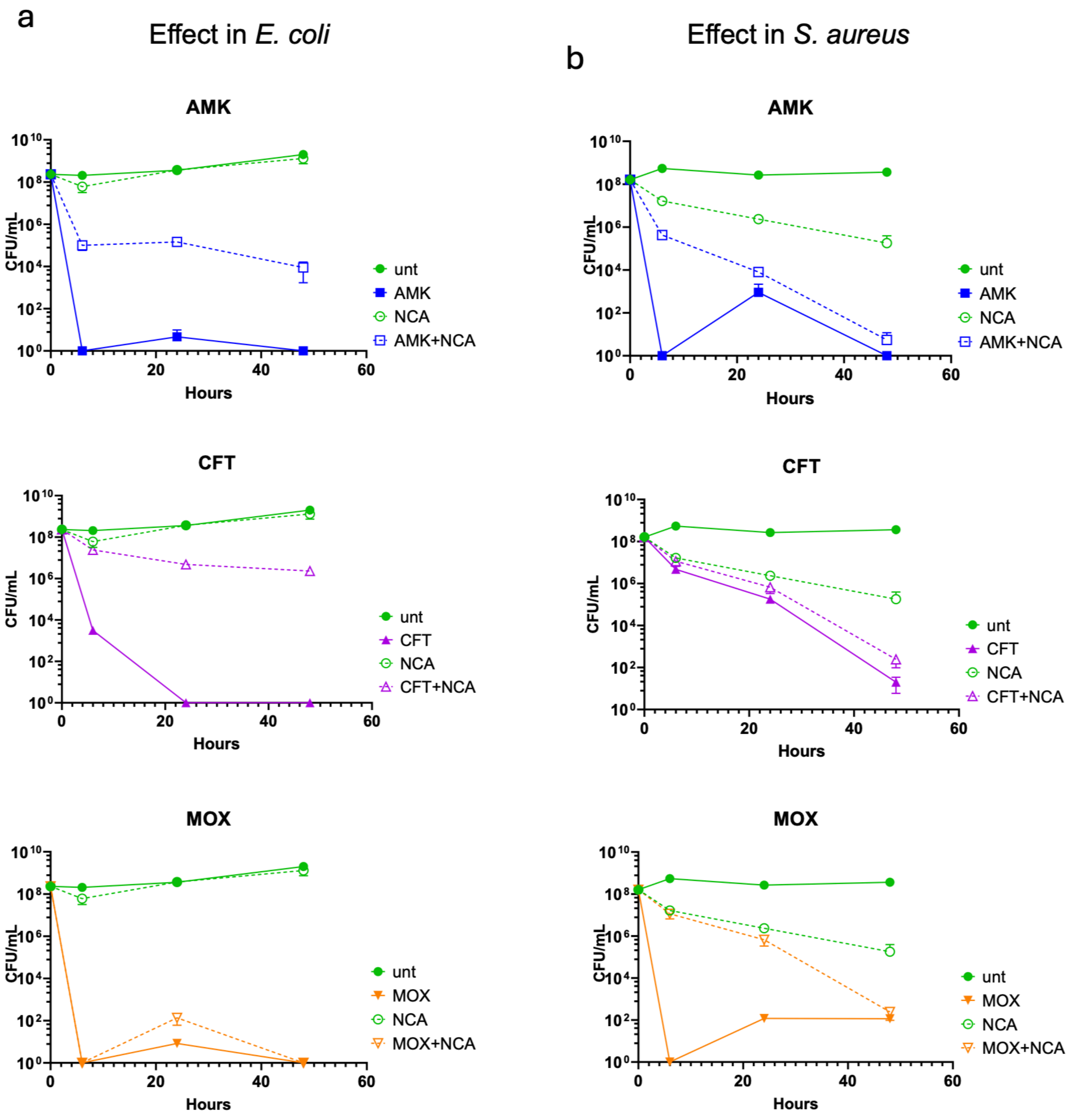


**Supplementary Figure 2. Effect of NCA on antibiotic-mediated killing in other bacterial species. (a)** *E. coli* and **(b)** *S. aureus* were exposed to NCA (10 µg/ml or 0.25 µg/ml, respectively) or left untreated, alongside 100× MIC of amikacin (AMK), ceftazidime (CFT), or moxifloxacin (MOX).

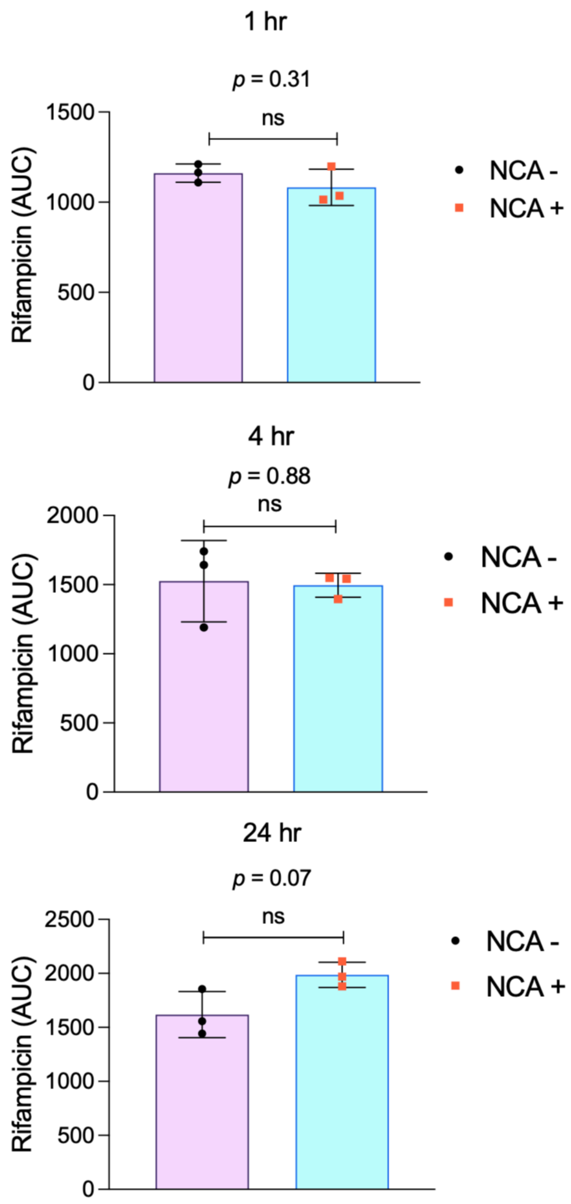


**Supplementary Figure 3. Effect of niclosamide (NCA) on intracellular rifampicin (RIF) concentrations in M. tuberculosis H37Rv.** Cells were treated with RIF (1 µg/ml) with or without NCA (2× MIC) for 1, 4, and 24 hours. Following treatment, cells were pelleted, washed twice with PBS, lysed by bead beating, and filter sterilized. Intracellular RIF levels were quantified by mass spectrometry. Asterisks indicate statistical significance; *p*-values were determined using a two-tailed Welch's *t*-test (n=3).

**Supplementary Table 2:** Log2 fold changes of commonly regulated DEGs that were significantly altered ($p < 0.02$) by at least 2-fold in response to the tolerance-promoting inhibitors (NCA, CCCP, and NIG). Corresponding values from the VAL group are included for comparison, along with the average Log2 FC difference.

| GENE_ID | Log2FC | | | | Average Log2 FC difference relative to VAL |
|---|---|---|---|---|---|
| | NCA | CCCP | NIG | VAL | |
| Rv3290c | 3.91 | 2.58 | 3.28 | 0.72 | 2.53 |
| Rv3289c | 3.64 | 2.33 | 3.08 | 0.54 | 2.48 |
| Rv1057 | 2.80 | 1.05 | 1.52 | -0.87 | 2.66 |
| MT2165.1 | 2.43 | 1.28 | 2.97 | -0.91 | 3.14 |
| Rv2428 | 2.31 | 1.30 | 2.42 | 0.69 | 1.32 |
| Rv2107 | 2.27 | 1.16 | 3.29 | -0.08 | 2.32 |
| Rv2429 | 2.21 | 1.26 | 1.99 | 1.60 | 0.22 |
| Rv0211 | 1.74 | 1.21 | 1.26 | 0.83 | 0.57 |
| Rv1286 | 1.58 | 1.22 | 1.48 | 0.85 | 0.58 |
| Rv1285 | 1.38 | 1.04 | 1.57 | 1.58 | -0.25 |
| Rv0059 | -1.49 | -1.00 | -1.17 | 0.36 | -1.58 |
| Rv0241c | -1.77 | -1.08 | -1.73 | 0.81 | -2.34 |
| Rv0242c | -1.83 | -1.05 | -1.84 | 0.98 | -2.55 |
| Rv0057 | -1.95 | -1.15 | -1.17 | 2.91 | -4.33 |
| Rv2988c | -2.15 | -1.43 | -1.02 | 1.27 | -2.80 |
| Rv2987c | -2.46 | -1.10 | -1.01 | 2.15 | -3.68 |

**Supplementary Table 3.** PAM sequences, sgRNA sequences, and forward/reverse primer sequences for CRISPRi knockdown of selected genes.

| Gene ID | pam Sequence | SgRNA target sequence | Forward primer | Reverse primer |
|---|---|---|---|---|
| Rv3290c | CGAGAAC | ATCAGCCCGCGGC CGCGCGGAT | GGGAATCAGCCCGC GGCCGCGCGGAT | AAACATCCGCGCGGC CGCGGGCTGAT |
| Rv3289c | GCAGAAA | GCACGCCATACCC GGTAGGTGC | GGGAGCACGCCATA CCCGGTAGGTGC | AAACACGCGAAGTCG AGGCCAC |
| Rv1057 | TGGGAAA | ATTGCGTGGTG GTGCCCGGCGCC | GGGAATTGCGTGG TGGTGCCCGGCGCC | AAACGGCGCCGGG CACCACCACGCAAT |
| Rv2428 | CTGGAAA | GGTCGTTGTGCTG TGCACGCCA | GGGAGGTCGTTGTG CTGTGCACGCCA | AAACGATCACACACG CCCCAAAGTT |
| Rv2107 | AAAGAAC | GGTACTGCTGCCC GTGGCTACT | GGGAGGTACTGCTG CCCGTGGCTACT | AAACCGAACTATTCG GCCGGCGTTTGC |
| Rv0059 | GTAGAAC | GCATGGGCGACC GCGCCGCAAT | GGGAGCATGGGCG ACCGCGCCGCAAT | AAACATTGCGGCGCG GTCGCCCATGC |
| Rv0241c | ATGGAAA | GCCGGTCACCAAC GACATCACCG | GGGAGCCGGTCACC AACGACATCACCG | AAACCGGTGATGTCGT TGGTGACCGGC |
| Rv0242c | CAGGAAT | ACGTAGGCCGACT TGGCCGACAG | GGGAACGTAGGCC GACTTGGCCGACAG | AAACCTGTCGGCCA AGTCGGCCTACGT |
| Rv0057 | CAAGAAT | AACGCGACGCCG TTTGTGTGGA | GGGAAACGCGACGCCG TTTGTGTGGA | AAACTCCACACAAA CGGCGTCGCGTT |
| Rv2988c | CAAGAAC | GTGGGTGCGTGCG GACGACCACG | GGGAGTGGGTGCGTG CGGACGACCACG | AAACCGTGGTCGTCC GCACGCACCCAC |
| Rv2987c | TCAGAAA | AAACCGGTTCGGGT GACCCGCT | GGGAAAACCGGTTCG GGTGACCCGCT | AAACAGCGGGTCACCCGA ACCGGTTT |
